# A new genome assembly of the pea cultivar ‘Caméor’ provides resources for functional genomics and genetics

**DOI:** 10.1101/2025.04.01.645976

**Authors:** Jonathan Kreplak, Petr Novak, Laura Ávila Robledillo, Grégoire Aubert, Baptiste Imbert, Parwinder Kaur, Quentin Gouil, Céline Lopez-Roques, Nathalie Rodde, Olivier Bouchez, Nadim Tayeh, Jiri Macas, Judith Burstin

## Abstract

Significant improvements in sequencing technologies have allowed the development of more contiguous genome assemblies in many plant species. The pea genome is characterized by its richness in repeated elements and its long and complex centromeres. This makes its assembly challenging. In this paper, we present an improved version of the genome sequence of the French cultivar ‘Caméor’. This sequence was obtained by combining Nanopore and PacBio long-read sequencing, Hi-C contact maps and Bionano maps. The assembly of centromeres was refined using a combination of FISH and ultra-long Nanopore read analyses. Overall, Cameor_v2 genome assembly is a highly continuous pea genome assembly with small total gap size and a large contig N50. In this version, the orientation of chromosomes was revised according to internationally accepted karyotype rules. Gene annotation statistics indicated a high completeness of gene sequences, with most gene sequences with 3’ and 5’ UTR. This genome assembly with its associated data constitute a useful resource for pea genetics, comparative mapping and functional genomics.

## Background & Summary

Pea (*Pisum sativum* L., 2n=14) is one of the most cultivated grain legumes in the world with soybean, common bean and chickpea^1^. Pea seeds, consumed both fresh (~21 M tons produced worldwide in 2022) and dry (14 M tons in 2022), are an essential source of proteins, fibres and micronutrients for human and animal nutrition. Moreover, the pea crop is an important component of agroecological cool-season cropping systems. Pea plants can acquire nitrogen through a symbiosis with N-fixing soil bacteria, reducing the need for synthetic fertilizers in crop rotations, and thus limiting greenhouse gas emission.

Pea breeding has long been hampered by the lack of a reference genome. The first draft assembly of the seven pea chromosomes was achieved for the French dry pea cultivar ‘Caméor’ using short-read sequencing in combination with physical and genetic mapping^2^. A significantly more contiguous assembly was produced for the Chinese variety ‘Zhongwan 6’ (ZW6)^3^ by using PacBio long-read sequencing. More recently, new pea genome assemblies were obtained for a vegetable Chinese pea variety, ‘Zhewan 1’ (ZW1)^4^, and the dwarf, wrinkled-seeds accession JI2822^5^.

The pea genome is rich in repetitive DNA consisting mainly of a diverse population of giant Ty3/gypsy Ogre elements^6^. In addition, pea chromosomes are characterized by highly elongated centromeres containing several separate domains of CENH3 chromatin (“metapolycentromeres”^7^). There are twelve families of satellite DNA associated with pea centromeric chromatin^8^, some of which form arrays of nearly identical monomers that are up to 2 Mb in size. These features make the full assembly of the pea genome, and especially its centromeres, a particularly challenging task. However, the use of ultra-long Oxford nanopore and PacBio HiFi reads in combination with cytogenetic mapping has recently led to a nearly complete assembly of the 177.6 Mb region of pea chromosome 6, including its 81.6 Mb centromere (CEN6), thus demonstrating the feasibility of this approach for assembling the complete pea genome^9^.

In the present study, we aimed to generate a high-quality chromosome-level assembly of *P. sativum* cv. ‘Caméor’ by combining newly generated PacBio HiFi and ultra-long Oxford Nanopore reads with Hi-C and optical mapping data. The centromeres of the remaining six pea chromosomes were integrated into chromosome-level pseudomolecules using the approaches developed for CEN6 assembly and characterized using CENH3 ChIP-seq data. This new genome assembly, with carefully annotated genes and repetitive DNA, constitutes a high-quality reference genome for pea. Gene expression and SNP data were linked to it in order to provide a comprehensive resource for research and breeding.

## Methods

### Genomic DNA sequencing

High molecular weight (HMW) DNA was prepared from the nuclei extracted from leaf tissues of pea cv. ‘Caméor’ seedlings, as described previously^10^. DNA quality was checked using field inversion gel electrophoresis (FIGE) to ensure that the DNA fragment size was >100 kb. Whole genome sequencing included 73.1 Gb PacBio HiFi reads and 119.6 Gb Nanopore reads ranging from 30 to 801 kb in length (N50 = 83.8 kb) that were generated as described in Macas et al. 2023^9^.

Additional Nanopore reads were obtained from leaf genomic DNA isolated using Qiagen Genomic-tips 100/G kit (QIAGEN, Hilden, Germany). At GeT-PlaGe (INRAE Toulouse), library preparation and sequencing were performed according to manufacturer’s instructions “1D gDNA selecting for long reads (SQK-LSK109)”. At each step, DNA was quantified using the Qubit dsDNA HS Assay Kit (Life Technologies). DNA quality was tested using a nanodrop spectrophotometer (Thermofisher) and size distribution and degradation was assessed using the Fragment analyzer (Agilent) DNF-464 HS Large Fragment Kit. Purification steps were performed using AMPure XP beads (Beckman Coulter). Several libraries were obtained. Libraries 1 and 2 were produced from 10 μg of DNA purified then sheared at 40kb using the megaruptor1 system (Diagenode). A size selection step using Short Read Eliminator XS Kit (Circulomics) was performed. The two libraries were loaded using SQK-LSK 109 kit (ONT) onto one Flongle (9fmol), two GridION (15fmol) and two PromethION (15fmol and 18fmol) R.9.4.1 flowcells. Library 3 was produced from 5 μg of DNA purified and then treated by an extra repair step with SMRTbell DAMAGE REPAIR KIT (PACBIO, 100-465-900). A size selection step using Short Read Eliminator XS Kit (Circulomics) was performed. The library was loaded using SQK-LSK 109 kit (ONT) onto one Flongle (8fmol) and one PromethION (15fmol) R.9.4.1 flowcells. Library 4 was produced from 20 μg of DNA purified and then treated by an extra repair step with SMRTbell DAMAGE REPAIR KIT (PACBIO, 100-465-900). A size selection step using Short Read Eliminator XL Kit (Circulomics) was performed. The library was loaded using SQK-LSK 109 kit (ONT) onto one GridION (15fmol) and one PromethION (15fmol) R.9.4.1 flowcells. In total, 249 Gb Nanopore reads were produced with N50 ranging from 15 to 22 kb.

At Walter and Eliza Hall Institute of Medical Research, two libraries were prepared, each using 2 μg high-molecular weight genomic DNA resuspended in 48 μl water, with the SQK-LSK109 kit according to the manufacturer’s instructions except for extended FFPE/End repair incubations (15 min. for each step instead of 5 min.). 600 ng and 1 μg of final libraries were recovered and loaded on two separate PromethION flow cells (R9.4.1 pore version). This generated 94 Gb Nanopore reads. Altogether, 462 Gb Nanopore reads and 73 Gb PacBioHiFi reads were used to construct the Cameor v2 genome assembly.

### High-throughput chromatin conformation capture (Hi-C)

To produce a high-quality chromosome level assembly, Hi-C maps were produced as described at https://www.dnazoo.org/methods. HiC sequencing was performed at GeT-PlaGe (INRAe Toulouse). Library quality was assessed using an Advanced Analytical Fragment Analyser and libraries were quantified by QPCR using the Kapa Library Quantification Kit. Sequencing has been performed on an Illumina HiSeq3000 using a paired-end read length of 2×150 pb with the Illumina HiSeq3000 Reagent Kits. Altogether, 780 million HiSeq3000 reads were produced representing 218 Gb total sequence.

### Optical maps

Optical maps were produced at CNRGV (https://cnrgv.toulouse.inrae.fr/fr) using the Bionano Irys system as described in Aury et al. 2022^11^. uHMW DNA was purified from 1g of dark treated young leaves according to the Bionano Prep Plant tissue DNA Isolation Liquid Nitrogen Grinding Protocol (30177 - Bionano Genomics) with the following specifications and modifications. Briefly, the leaves were fixed in fixation buffer containing formaldehyde, rinsed, manually chopped and then disrupt with rotor stator in homogenization buffer. Nuclei were washed and then embedded in agarose plugs. After overnight proteinase K digestion in the presence of Lysis Buffer and one hour treatment with RNAse A (Qiagen), plugs were washed three times in 1x Wash Buffer and three times in 1x TE Buffer (ThermoFisher Scientific). Then, plugs were melted two minutes at 70°C and solubilized with 2 μL of 0.5 U/μL AGARase enzyme (ThermoFisher Scientific) for 45 minutes at 43°C. A dialysis step was performed in 1x TE Buffer (ThermoFisher Scientific) for 45 minutes to purify DNA from any residues. The DNA samples were quantified using the Qubit dsDNA BR Assay (Invitrogen) and the presence of mega base size DNA was visualized using pulsed field gel electrophoresis (PFGE). Labeling and staining of the uHMW DNA were performed according to the Bionano Prep Direct Label and Stain (DLS) protocol (30206 - Bionano Genomics). Briefly, labelling was performed by incubating 750 ng genomic DNA with 1× DLE-1 Enzyme for 2 hours in the presence of 1× DL-Green (Bionano Genomics) and 1× DLE-1 Buffer. Following proteinase K digestion and DL-Green cleanup, the DNA backbone was stained by mixing the labeled DNA with DNA Stain solution (Bionano Genomics) in presence of 1× Flow Buffer and 1× DTT (Bionano Genomics), and incubating overnight at room temperature. The DLS DNA concentration was measured with the Qubit dsDNA HS Assay (Invitrogen) and was loaded on the Saphyr chip. Loading of the chip and running of the Bionano Genomics Saphyr System were all performed according to the Saphyr System User Guide(30247 - Bionano Genomics). Data processing was performed using the Bionano Genomics Access software https://bionanogenomics.com/support-page/bionano-access-software/). 1 152 Gb of filtered data (>150kb) were produced and assembled producing 391 genome maps with a N50 of 25 Mbp for a total genome map length of 3808 Mbp.

### Fluorescence in situ hybridization

The mitotic chromosomes used for FISH were prepared from synchronized root tip meristems as described^12^. Satellite repeats were detected with oligonucleotide probes designed according to their consensus sequences and labelled with 5’-biotin during synthesis (Integrated DNA Technologies, Leuven, Belgium). Alternatively, cloned fragments of satellite repeats obtained by PCR amplification with specific oligonucleotide primers and labelled by nick translation were used as FISH probes. The labelling and FISH were performed as described^8^, with hybridization and wash temperatures adjusted to account for AT/GC content and stringency of hybridization, allowing for 10–20% mismatches. Slides were counterstained with 4’,6-diamidino-2-phenylindole (DAPI) in Vectashield mounting medium (Vector Laboratories, Burlingame, CA, USA) and examined using a Zeiss AxioImager.Z2 microscope with an Axiocam 506 monocamera. Images were captured and processed using ZEN 3.2 software (Carl Zeiss GmbH).

### ChIP-seq detection of centromeric chromatin

The positions of the centromeres in the assembled pseudomolecules were determined by mapping the ChIP-seq reads generated for the two variants of pea CENH3 proteins, CENH3-1 and CENH3-2^9,12^. The ChIP-seq experiments were performed with native chromatin^7^ using custom antibodies. DNA fragments isolated from the immunoprecipitated samples were sequenced together with the corresponding control samples (input; digested chromatin not subjected to immunoprecipitation) on the Illumina platform (Admera Health, NJ, USA) in paired-end, 150 bp mode. Duplicate experiments, including independent chromatin preparations, were performed for each CENH3 variant with either one antibody (P23 for CENH3-2 targeting the epitope “TPRHARENQERKKRRNKPGC”) or with two different antibodies (P22 and P43 for CENH3-1 targeting the same epitope “GRVKHFPSPSKPAASDNLGKKKRRCKPGTKC” but raised in rabbit and chicken, respectively).

The resulting reads were quality filtered and trimmed using Trimmomatic^13^ (minimum allowed length = 100 nt), yielding 122–211 million reads per sample, which were mapped to the assembly using Bowtie 2 version 2.4.2^14^ with the options -p 64 -U. The subsequent analysis was performed with the complete output of the Bowtie 2 program and with the output in which all multimapped reads were filtered out. The filtering of multimapped reads was performed with Sambamba version 0.8.1^15^ with the options “-F [XS] == null and not unmapped and not duplicate”. Regions with statistically significant ChIP/input enrichment ratios were identified by comparing ChIP- and input-mapped reads using the program epic2^16^ and the parameter “--bin-size 200”. An alternative identification of enrichment was performed using MACS2^17^ version 2.1.1.20160309 with the default settings. The ChIP/input ratio was calculated for plotting purposes using bamCompare (version 3.5.1) from the deepTools package ^18^. The program was run with the parameter “-binSize 200” to calculate the log2 ratio for the window size of 200 nt. The resulting data was recorded using the rtracklayer package from R^19^. The complete Illumina reads preprocessing pipeline and ChIP-seq analysis pipeline were implemented as a Snakemake workflow and executed using a Singularity container. The source code for these pipelines is available at https://doi.org/10.5281/zenodo.14802196 and https://doi.org/10.5281/zenodo.14801796.

### Chromosome-scale genome assembly

Nanopore reads were trimmed using porechop^20^ and filtered with NanoFilt^21^ to keep only those with a quality of 8 and a length of at least 10 kb. Contigs were assembled using flye (Kolmogorov et al. 2019) and then scaffolded using Hi-C data with the Juicer^22^ and 3D-DNA pipeline^23^. The assembly was refined through a round of manual corrections with JuiceBox Assembly Tools^24^. Genetic maps^25,26^ were then anchored using^27^ to assess chromosome sequences. The assembled pseudo-molecules were corrected first by two rounds of Racon^28^ using nanopore long reads and one round of medaka (“GitHub - Nanoporetech/Medaka: Sequence Correction Provided by ONT Research” n.d.). Then, the genome sequence was polished by one round of Polca^29^ with short-reads and one with HiFi long reads. Centromeres were manually corrected^9^ using FISH data. Finally, assembly gaps were filled using HiFi sequence reads.

### Repetitive DNA annotation and masking

Tandem repeats and satellites were annotated using TideCluster v.1.6 (https://doi.org/10.5281/zenodo.7885625), a wrapper for TideHunter^30^. Satellite repeats with a monomer size ranging from 40 bp to 3 kb and a minimum array length of 5 kb were annotated using the default TideCluster settings. Satellites with a monomer size between 10 to 39 bp and a minimum array length of 5 kbp were identified using TideCluster with parameters -T “-p 10 -P 39 -c 5 -e 0.25” -m 5000.

LTR retrotransposons (LTR-RT) were annotated using DANTE v0.2.5 and the DANTE_LTR v0.4.0.4 pipeline^31^ on the RepeatExplorer Galaxy server^32^. The sequences of the identified LTR-RT elements were used to create a custom library of LTR-RT elements using the “dante_ltr_to_library” script from the DANTE_LTR repository (https://doi.org/10.5281/zenodo.7891007).

A custom library of Class II transposable elements was obtained from previously published datasets^33,34^ and using RepeatExplorer clustering procedure 1^35^ on unassembled Illumina paired-end reads. Contigs corresponding to Class II retrotransposons with a minimum read depth of 5 reads and a minimum length of 100 bp were obtained using tools on the RepeatExplorer Galaxy server. A custom library of LINE elements was created by extracting regions with LINE protein coding domains identified by DANTE, along with the upstream and downstream 4kb flanking regions. The extracted genomic sequences were split into 100 nt fragments and analyzed by RepeatExplorer clustering. Contigs corresponding to LINE elements with a read depth of at least 3 reads and a minimum length of 150 nt were converted into a custom library. Consensus sequences of rRNA gene arrays including intergenic spacer sequences were fully reconstructed from the RepeatExplorer contigs.

All custom libraries were concatenated and used as a library for RepeatMasker^36^ search. The RepeatMasker (v 4.1.5) search was performed with options “-xsmall -no_is -e ncbi”. All regions annotated as mobile elements with RepeatMasker based on custom library search which overlapped with satellite repeats annotated by TideCluster were removed from the annotation using bedtools (v 2.31.1) with command “bedtools subtract”.

The resulting GFF3 was then merged with the DANTE annotation using a custom R script. The classification of mobile elements in the annotation files corresponds to the classification system used in the REXdb database^37^.

For the final repeat-masking process, all of the above repeat annotation GFF3 files were consolidated. We merged the annotated regions into a single BED file using the bedtools merge tool^38^.

The complete repeat annotation pipeline was implemented as a Snakemake workflow and executed using a Singularity container. The source code for the pipeline is available at https://doi.org/10.5281/zenodo.14801742.

### Gene annotation

Several RNA-seq datasets were mapped onto the Cameor v2 assembly using STAR^39^ with the following options: --outSAMstrandField intronMotif --twopassMode Basic -- outFilterMultimapNmax 20 --alignSJDBoverhangMin 1. Datasets correspond to different plant tissues produced under control conditions^40^ (PRJNA267198); to developing seed tissues under water and sulfur stress^41^ (PRJNA517587); to plant tissues produced under cold-stress^42^ (PRJNA543764). Only reliable junctions for each alignment files were kept using Portcullis^43^. Stringtie2^44^ was then used to assemble a transcriptome for each condition, merging replicates when this was possible. Then, Mikado^45^ selected the best transcript for each dataset, yielding four different assemblies.

Transcriptomes assemblies and protein sets from different legumes (*V. faba, L. culinaris, M. truncatula, L. japonicus)* were used as hints to run the Eukaryote EuGene pipeline version 1.6.5 ^46^. Helixer^47^ was run on the genome using the land_plant 0,3 model and general recommendations for plants (subsequence-length at 64152, overlap-offset at 32076 and overlap-core-length at 48114). Both annotations were then merged using gffcompare^48^ and functionally annotated using emapper on eggNOG 5,0^49^. Only gene models with less than 50% of their coding sequences overlapping a transposable element and not functionally annotated as a repeat were kept. Terms used for this functional annotation screening were: transposon|LTR|copia|GAG|helicase|integrase|gypsy|transposition|ribonuclease|transcrip tase|polymerases|mobile. The distribution of genes and repeated elements across the genome is represented on Figure 1.

**Figure 1:**
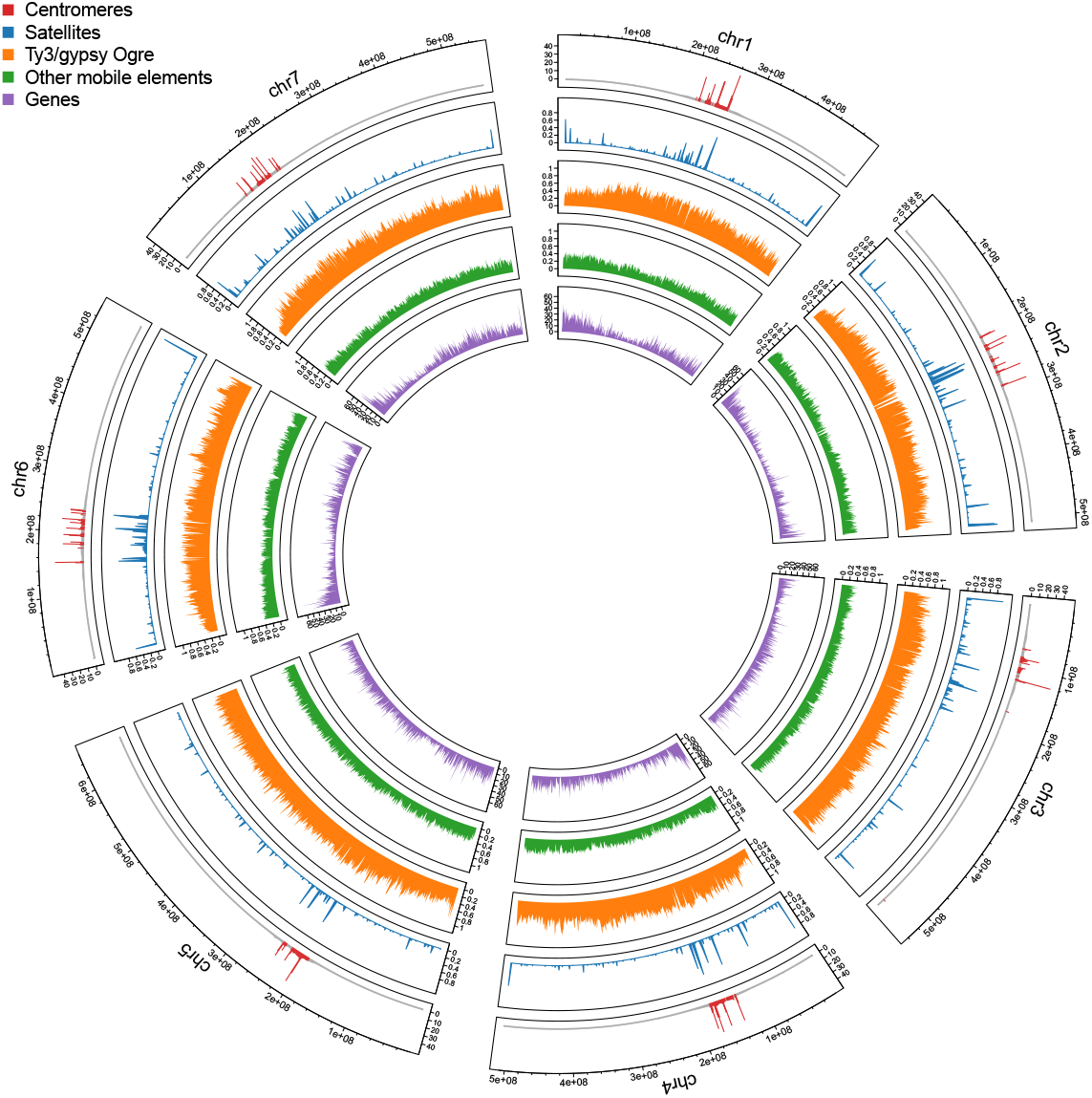
Distribution of genomic features on the chromosomes of *P. sativum* cv. ‘Caméor’. The tracks on the Circos plot are arranged from outside to inside, and represent: (1) position of centromeres as determined by ChIP-seq with the CENH3 antibody, shown as ChIP/input ratio; (2) density of satellite DNA; (3) density of Ty3/gypsy Ogre elements; (4) density of other mobile elements; (5) number of protein-coding genes. All densities, gene numbers and ChIP/input ratios are averaged over 1 Mb windows.

### Centromeric regions

Due to the large size and repeat complexity of the pea centromeres, these regions were subjected to additional rounds of review and possible correction using the following procedure: (i) the order and orientation of contigs was verified by the presence of ultra-long nanopore reads spanning the gaps between adjacent contigs, (ii) when possible, incomplete arrays of satellite repeats were extended or fully included using contigs generated by assembling HiFi reads as described in Macas et al. 2023^9^, (iii) the presence and arrangement of specific satellite repeats was verified by fluorescence in situ hybridization on mitotic chromosomes as in Macas et al. 2023^9^.

The centromere structure was analyzed by combining CENH3 ChIP-seq data with repeat and gene annotations. The centromere positions on the assembled pseudomolecules were defined as regions bounded by the outermost CENH3 domains, which also define the extent of primary constriction on metaphase chromosomes (Figure 1). The size of these regions ranged from 35.8 Mb (CEN5) to 83.9 Mb (CEN2) (Table 1). The CENH3 chromatin domains were mostly located on the arrays of satDNA and there were eleven different satDNA families associated with CENH3. In addition, there were arrays of CENH3-free satellites in all centromeres. The distribution patterns of satDNA were unique for each centromere (Figure 2).

**Table 1:**
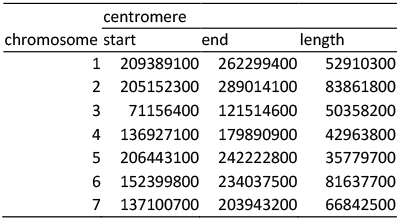
Coordinates of the centromeres of the seven chromosomes of Cameor_v2 genome assembly.

**Figure 2:**
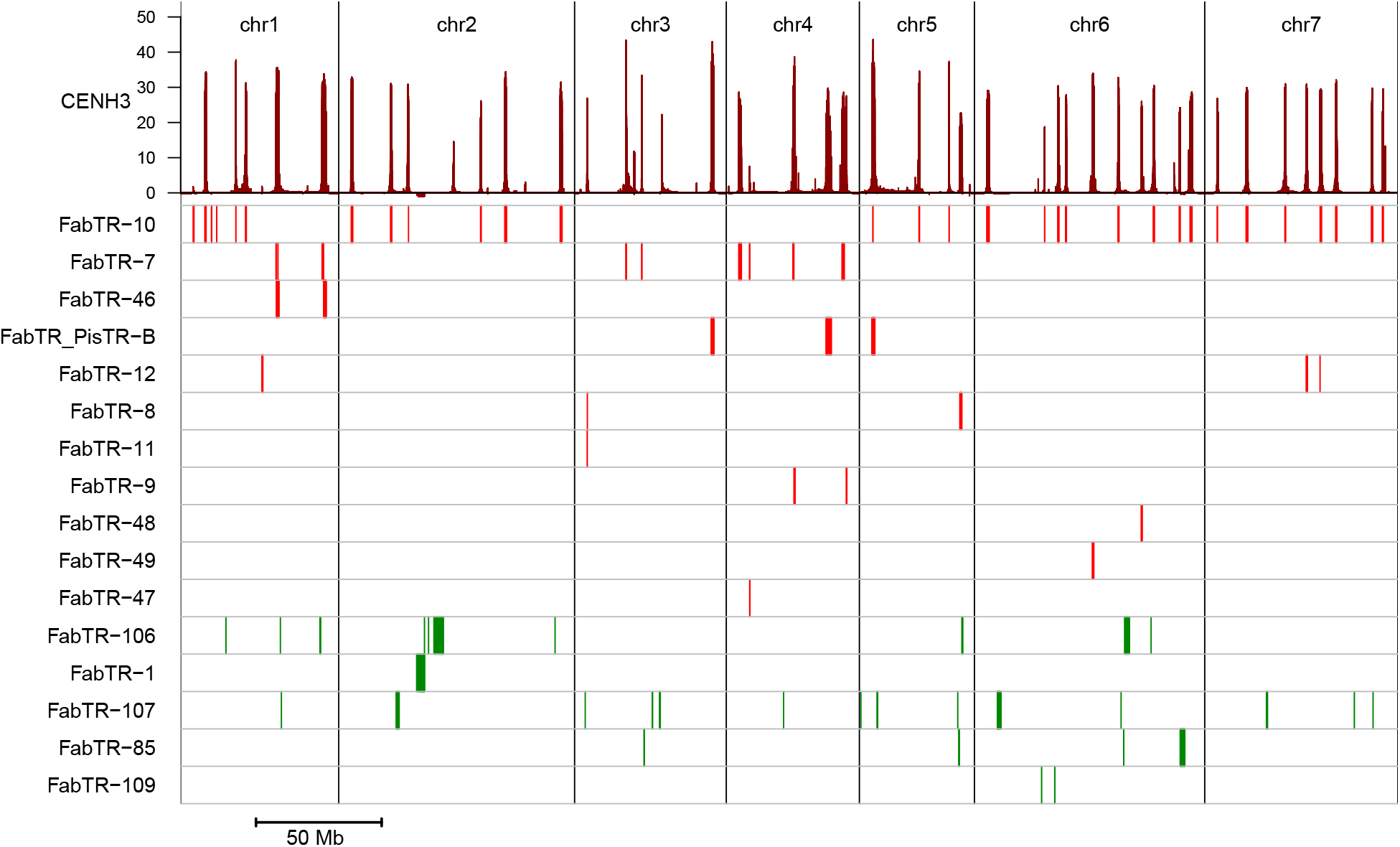
Structure of the centromeres. The upper panel shows the distribution of multiple centromere domains along the centromeric regions (see Table 1 for the coordinates of the plotted regions). The domains are revealed by the increased ChIP/input read ratio in ChIP-seq with the CENH3 antibody. The bottom panel shows the distribution of sixteen families of centromeric repeats, with each row corresponding to a different repeat family. The satellites highlighted in red are associated with CENH3 chromatin.

### Resources for genetics, comparative and functional genomics

RNA-seq datasets used for annotation were mapped onto the Cameor v2 assembly. Moreover, the SNP context sequences of genotyping arrays were mapped onto this genome version: the Infinium GENOPEA array^25^ and the Axiom array^26^. The set of exome capture SNP^50^ was recalled on the genome. Briefly sequenced reads were trimmed for adaptor sequence using cutadapt 1.8.3 ^51^. The GATK variant calling best practices pipeline was run on the new version exome content using elprep 5.1.3^52^, and GATK 4.2.6.1^53^ was used to merge and produce a SNP VCF file. Furthermore, Cameor_v2 was integrated in the OrthoLegKB database^54^in order to enable comparative genomics and translational approaches using this new genome version. Syntenic relationships between Cameor_v2 and other legumes’ genomes was investigated as described in Imbert et al 2023 ^54^.

## Data Records

Raw sequence data are available from the European Nucleotide Archive (https://www.ebi.ac.uk/ena/browser/home) study PRJEB54858 under accession numbers ERR9972527-ERR9972530 (PacBio HiFi), ERR9980778-ERR9980797 (Oxford Nanopore), and ERR9981080-ERR9981086 (CENH3 ChIP-seq). The genome sequence and the HiC sequencing data are available under the study PRJEB78861. The genome sequence and associated data are also available at : https://doi.org/10.57745/5MDE99.

## Technical Validation

Cameor_v2 final assembly includes seven pseudomolecules and 1,387 unplaced scaffolds. Unplaced sequences account for 3% of the total assembly length (120 Mb). Contigs assembly based on Nanopore long-reads, Hi-C scaffolding and a careful manual curation of centromeric regions, led to a significantly more continuous genome assembly of *P. sativum*, with smaller total gap size in the assembly and the larger contig N50 (Table 2). A manual validation of the assembly was done using bionano maps. Two super-scaffolds were oriented based on this curation. Additionally, the assembly of centromeric regions was verified by visualization of specific families of satellite DNA using FISH. The comparison of Cameor_v2 to Cameor_v1 confirmed that this genome assembly was significantly improved, especially for centromeric regions (Figure 3).

**Table 2:**
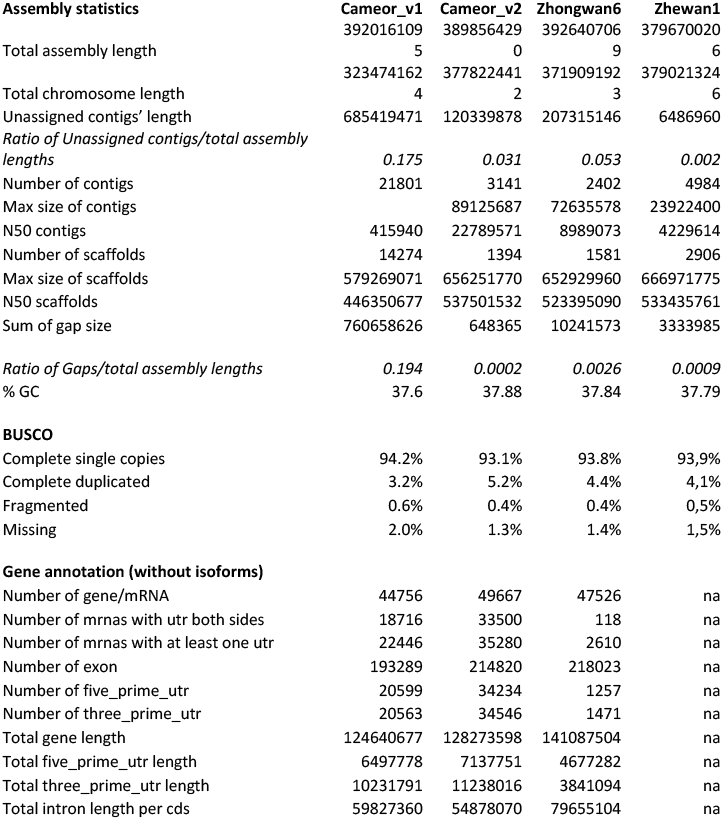
Assembly statistics of Cameor_v2 and published pea genome sequences of Cameor_v1 ^2^, ZW6 ^3^ and ZW1 ^4^.

**Figure 3:**
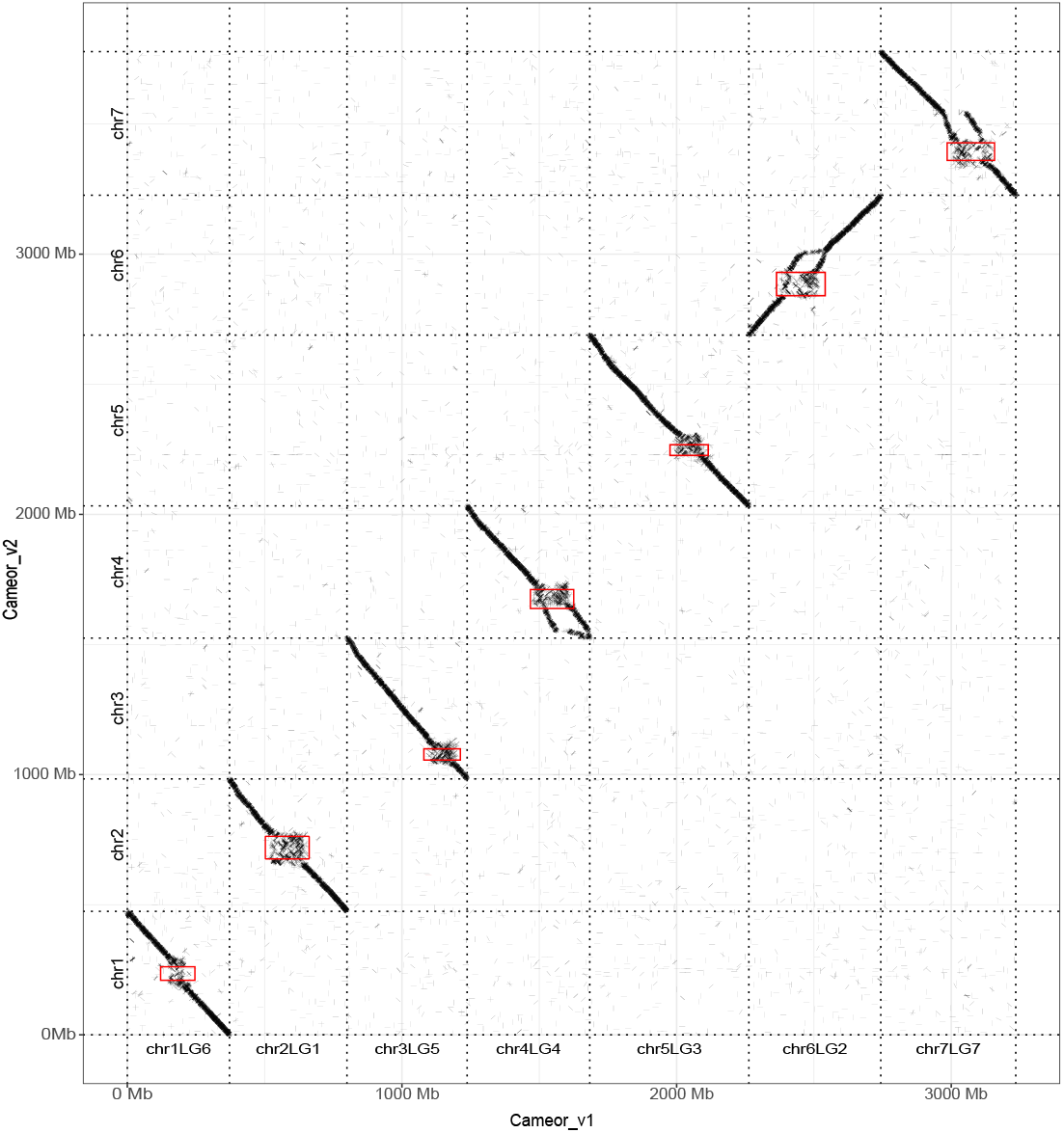
Dot-plot comparing the Cameor_v2 genome sequence with the published pea genome sequence Cameor_v1(^2^); Dotted red squares indicate the limits of centromeres in Cameor_v2 assembly.

Thanks to the refined assembly of centromeres in Cameor_v2, the orientation of chromosomes was revised according to the internationally accepted karyotype rules, with the short arm being placed at the top, and the long arm at the bottom of chromosomes. Chromosome sequences of Cameor_v2 thus start from the telomere of the short arm, through the centromere, to the telomere of the long arm at the end of the pseudomolecule. This led to the inversion of chromosomes 1, 2, 3, 4, 5 and 7 in Cameor_v2 as compared to Cameor_v1 (Figure 3).

The quality of gene annotation was compared with other pea genome assemblies published recently. A BUSCO^55^ analysis showed that Cameor_v2 has more complete duplicate and slightly less missing genes than other published pea genome assemblies (Table 2; Kreplak et al. 2019; Yang et al. 2022; Liu et al. 2024), probably thanks to an increased completion of gene detection and sequences. Interestingly, the number of genes with both 3’ and 5’ UTR was higher than in other genome assemblies, indicating a better completeness of genes.

Comparing published pea genome sequences (Figure 4) highlighted high collinearity between Cameor_v2 and genome assemblies of ZW6 and ZW1. Some breaks in collinearity occur, mostly near centromeres or telomeres (Figure 4): one on Chr4 and Chr6, three on Chr7 in comparison with ZW6; one on Chr2 and 4, two on Chr1 and Chr3, three on Chr5, and four on Chr7 in comparison with ZW1. Some more complex patterns were identified with ZW1. These breaks in collinearity may be due to real rearrangements (inversions, duplications) in the genomes of ‘Caméor’, ‘Zhongwan 6’ and ‘Zhewan 1’; or may be due to assembly artifacts, in regions that are difficult to assemble due to repetitive elements.

**Figure 4:**
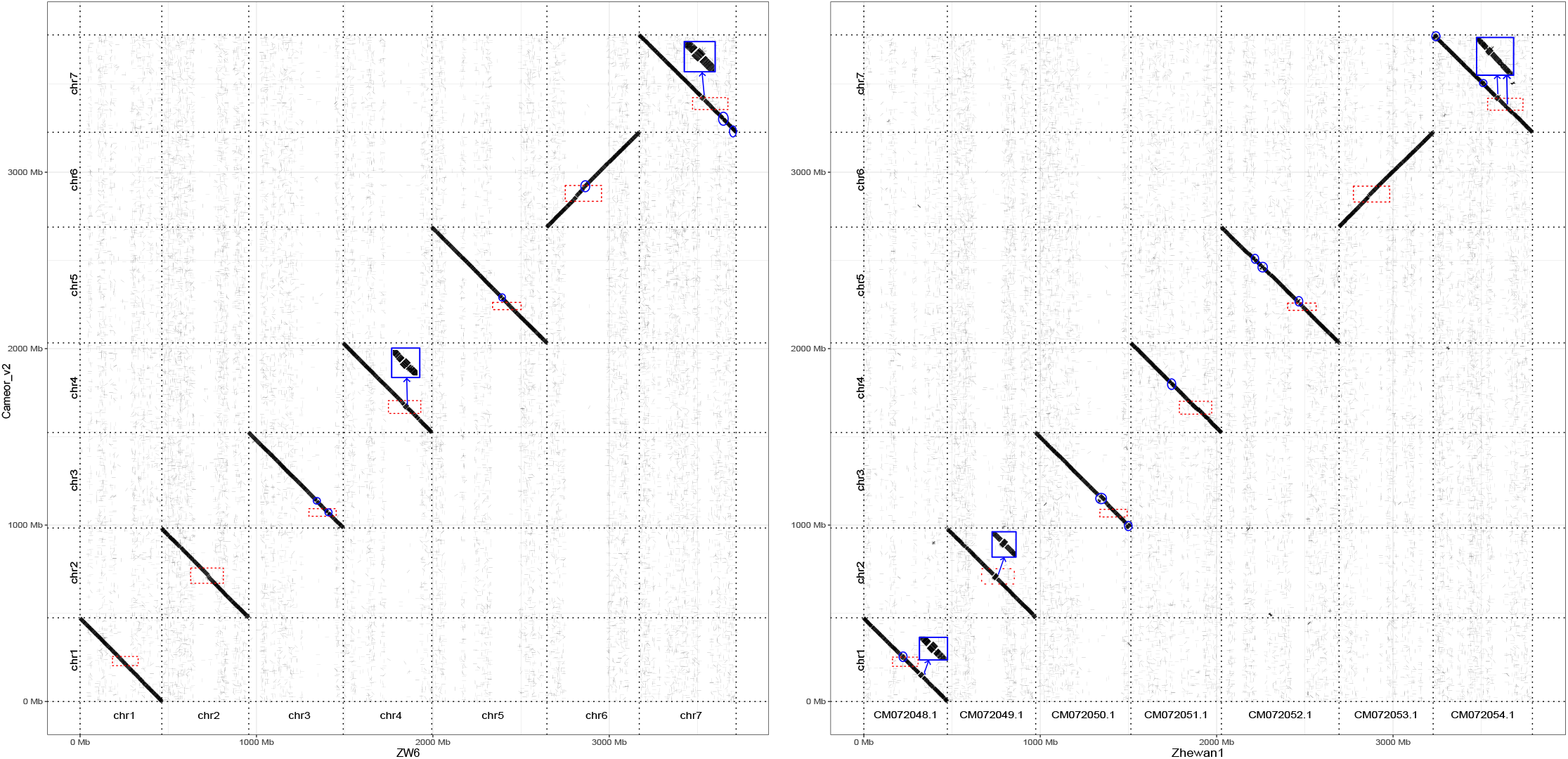
Dot-plot comparing the Cameor_v2 genome sequence with the published pea genome sequences of ZW6 ^3^ and ZW1 ^4^; Dotted red squares indicate the limits of centromeres in Cameor_v2 assembly; blue squares and circles highlight breaks in collinearity.

Assessing synteny and collinearity conservation between Cameor_v2 and other legume genomes showed that improving the quality of the genome sequence of ‘Caméor’ helped to detect and refine the syntenic relationships with other legume genomes (faba bean, lentil, chickpea, and the model species *M. truncatula* Gaertn., Figure 5). Larger syntenic blocks were identified for all chromosomes and gaps in centromeric regions were reduced in length (Figure 5). Syntenic blocks, i.e. conserved chains of orthologous genes across chromosome pairs, were fewer, but longer and included more genes when Cameor_v2 was used, as compared to Cameor_v1, certainly due to improved gene annotation and gene order (Table 3).

**Table 3:** Statistics of syntenic relationships between the pea genome assemblies Cameor_v1 and Cameor_v2 and genome assemblies from faba bean ^56^, lentil ^57^, chickpea ^58^, and *M. truncatula* Gaertn ^59^. Only orthologous genes found in the Cameor genome were considered. OrthoFinder^60^ was employed to identify orthogroups, using the genome of mung bean (*Vigna radiata* L.)^61^ as an outgroup. Orthogroups served as anchors to define syntenic blocks among the genome sequences, using MCScanx ^62^. MCScanX was run with its default parameters, except for the minimum number of genes required to define syntenic blocks, which was set to 10.

| Statistics | Cameor_v2 | Cameor_v1 |
| --- | --- | --- |
| Total number of protein-coding genes | 49 667 | 44 958 |
| Number of genes with orthologs | 33 072 | 30 439 |
| Number of orthologous collinear genes | 20 457 | 12 771 |
| % of collinear genes among orthologs | 62 | 42 |
| Mean orthologous gene count per block | 96 | 46 |
| Number of syntenic blocks | 703 | 814 |
| Mean block size | 18 471 838 | 11 063 388 |

**Figure 5:**
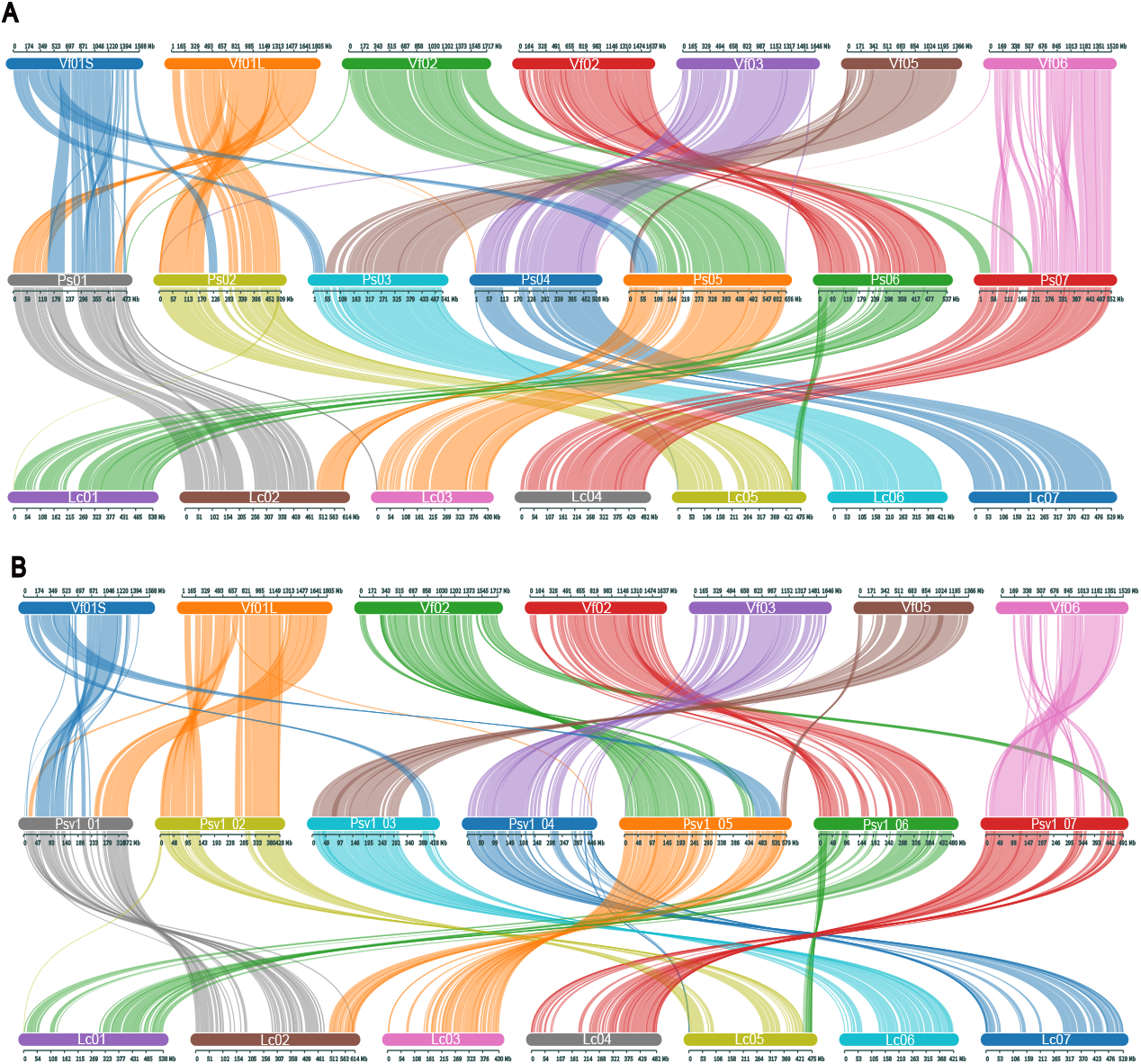
Synteny between chromosomes of *Pisum sativum* (Ps01 to Ps07), *Vicia faba* (Vf01 to Vf06)^56^, *Lens culinaris* (Lc01 to Lc07)^57^. A: using *P. sativum* Cameor_v2 genome; B: using *P. sativum* Cameor_v1 genome. Synteny was visualized using SynVisio^63^.

## Usage Notes

Genomic browser (JBrowse) including assembly annotation and ChIP-seq data tracks can be accessed from http://w3lamc.umbr.cas.cz/lamc/?page_id=8

## Code Availability

Repeat Annotation: https://doi.org/10.5281/zenodo.14801742 Illumina preprocessing: https://doi.org/10.5281/zenodo.14802196 ChiP-Seq analysis: https://doi.org/10.5281/zenodo.14801796

## Acknowledgements

This work was performed in collaboration with the GeT core facility, Toulouse, France (GeT, https://doi.org/10.15454/1.5572370921303193E12). GeT core facility was supported by France Génomique National infrastructure, funded as part of “Investissement d’avenir” program managed by Agence Nationale pour la Recherche (contract ANR-10-INBS-09). Nanopore sequencing and bionano maps done at INRAE were financed by the Project Investissements d’Avenir PeaMUST under the grant number ANR-11-BTBR-0002. JM, PN and LAR were supported by the Czech Science Foundation grant 24-10036S. Computational resources and data storage facilities were in part provided by the ELIXIR-CZ Research Infrastructure Project (LM2023055).

## Author contributions

JK produced the genome assembly and annotated the genes; GA managed the production of plant tissues and DNA extractions; BI and NT analysed the conservation of synteny; JM, PN and LAR produced ultra-long Nanopore reads and performed the FISH and ChIP-seq analysis, corrected the assembly, especially the centromeric and repeated regions, and annotated repeat elements; PK managed the production of HiC libraries at UWA; QG managed the production of Nanopore reads at WEHI; CLR managed the production of Nanopore reads at Get-Plage; NR managed the production of Bionano maps at GENOTOUL; OB managed the sequencing of HiC libraries. JB coordinated the project; JB and JM drafted the manuscript. JK, GA, PN, QG, CLR, NR, OB contributed to writing the manuscript. All authors read the manuscript.

